# RSAT matrix-clustering: dynamic exploration and redundancy reduction of transcription factor binding motif collections

**DOI:** 10.1101/065565

**Authors:** Jaime Abraham Castro-Mondragon, Sébastien Jaeger, Denis Thieffry, Morgane Thomas-Chollier, Jacques van Helden

## Abstract

Transcription Factor (TF) databases contain multitudes of motifs from various sources, from which non-redundant collections are derived by manual curation. The advent of high-throughput methods stimulated the production of novel collections with increasing numbers of motifs. Meta-databases, built by merging these collections, contain redundant versions, because available tools are not suited to automatically identify and explore biologically relevant clusters among thousands of motifs. Motif discovery from genome-scale data sets (e.g. ChIP-seq peaks) also produces redundant motifs, hampering the interpretation of results. We present *matrix-clustering*, a versatile tool that clusters similar TFBMs into multiple trees, and automatically creates non-redundant collections of motifs. A feature unique to *matrix-clustering* is its dynamic visualisation of aligned TFBMs, and its capability to simultaneously treat multiple collections from various sources. We demonstrate that *matrix-clustering* considerably simplifies the interpretation of combined results from multiple motif discovery tools and highlights biologically relevant variations of similar motifs. By clustering 24 entire databases (>7,500 motifs), we show that *matrix-clustering* correctly groups motifs belonging to the same TF families, and can drastically reduce motif redundancy. *matrix-clustering* is integrated within the RSAT suite (http://rsat.eu/), accessible through a user-friendly web interface or command-line for its integration in pipelines.

## INTRODUCTION

Transcription Factor Binding Motifs (TFBM) – simply called *motifs* below – are models describing the binding specificity of a transcription factor (TF). Such motifs are generally obtained by aligning the sequences of several binding sites, and summarizing the nucleotide frequencies per position. Motifs are commonly represented as position-specific scoring matrices (PSSMs) (1) and visualized as sequence logos (2). Although the adequacy of PSSMs has been questioned for some particular TF classes (3–6), e.g. in cases of dependencies between adjacent nucleotides, they are still the most widely used method to represent the binding specificity of a TF. Thousands of PSSMs are available in private or public databases, such as JASPAR (7), TRANSFAC (8), Cis-BP (9), FootprintDB (10), HOCOMOCO (11), which constitute key resources to interpret functional genomics results. A well-known issue with these databases is motif redundancy (12), caused by various reasons: (i) for a given TF, multiple PSSMs can be built from different collections of sites characterized with alternative methods (i.e. DNase-Seq, SELEX, Protein-Binding Microarrays (PBMs), ChIP-seq, etc); (ii) the binding specificity is often conserved between TFs of the same family; (iii) some databases contain PSSMs obtained from orthologous TFs in different organisms; (iv) some unrelated TFs recognize similar DNA motifs.

In addition to this intra-database redundancy, inter-database redundancy and the exponential growth of motif collections are becoming a major issue. Indeed, the development of high-throughput methods to characterize genome-wise TF binding locations (e.g. ChIP-seq, ChIP-exo) has led to an explosion of motifs, with a fast expansion of databases (e.g. JASPAR 2016 almost doubled in size since its 2014 version, from 590 to 1092 motifs) (12). In parallel, recent studies targeting many TFs (13, 14) resulted in collections with as many motifs as reference databases. This constant increase in the number of motifs and redundant collections represents a real challenge for the community. Which collection to use? How important is the overlap between the different collections? Efforts to collect and integrate numerous up-to-date collections into a single metadatabase like FootprintDB (10) or Cis-BP (9) are critical for the community. These metadatabases however do not deal yet with the redundancy issue, and keep increasing in size. This now constitutes a bottleneck, by drastically increasing the time needed to compare motifs or to scan sequences with a complete motif database.

Analysis of high-throughput datasets (e.g., from ChIP-seq experiments) also produces sets of redundant motifs. It is common practice to simultaneously use multiple *de novo* motif discovery tools (15–18), in order to benefit from their complementarity. While some motifs will be discovered exclusively by a given tool, most will be found independently by different tools, hence producing redundant motifs with small variations in length and/or nucleotide frequencies at some positions. Such variations may be important biologically, but remain undetected when inspecting unordered collections of motif logos.

Motif redundancy can be automatically reduced by identifying sets of similar motifs and clustering them. Quantifying the similarity between motifs is nevertheless far from trivial. Many efforts have been done to develop statistical methods and to find adequate comparison metrics between motifs, each one with its own strengths and drawbacks (19–36). Despite this intensive research activity to refine motif similarity metrics, no general consensus has emerged about the best one. Currently, a handful of tools are available for motif comparison: *STAMP (22, 37), TomTom (23), MATLIGN (26), macro-ape* (27), *DMINDA (35), DbcorrDB (34) and* RSAT *compare-matrices (38)*. Other tools are specialized in motif clustering: *STAMP (22), m2match (25), MATLIGN* (26), *GMACS (28), DMINDA (35)* and *motIV (Bioconductor package)* (see Table 1 for a comparison of their capabilities). However, each of these tools presents some limitations: analysis based on a single metric, restricted number of input motifs, static visualisation interfaces.

**Table 1.** **Features of software tools available to perform clustering of PSSMs.**

**Figure 5**
|  | Matlign | STAMP | m2match | DMINDA | motIV | GMACS | matrix-clustering |
| --- | --- | --- | --- | --- | --- | --- | --- |
| Year | 2007 | 2007 | 2013 | 2014 | 2014 | 2015 | 2016 |
| Clustering method |  |  |  |  |  |  |  |
| Hierarchical | YES | YES | YES | no | YES | no | YES |
| SOTA | no | YES | no | no | YES | no | no |
| Genetic Algorithm | no | no | no | no | no | YES | no |
| minimum-spanning-tree | no | no | no | YES | no | no | no |
| Multiple alignment of logos | no | no | YES | no | no | no | YES |
| Alignment with internal gaps | no | YES | no | no | no | no | no |
| Logo tree | no | no | YES | no | no | no | YES |
| Familial binding profiles | YES | YES | YES | no | YES | no | YES |
| Partitioning of input motif set in distinct clusters | no | YES | YES | no | no | YES | YES |
| Consensus/Logo at tree root | no | YES | YES | no | YES | no | YES |
| Consensus/Logo at each branch | no | no | no | no | no | no | YES |
| Multiple collections as input | no | no | no | no | no | no | YES |
| Accessible via website | YES | YES | YES | YES | no | no | YES |
| Restriction on number of matrices | no | no | 30 (trial account) | no | no | no | no |
| Publication | PMID: 17559640 | PMID: 17478497 | PMID: 23555204 | PMID: 24753419 | Bioconductor package | PMID: 25627106 | This article |

We have developed *matrix-clustering* within the RSAT suite (39), motivated by the crucial need for a tool to cluster similar motifs, align them to facilitate visual comparison, explore each cluster in a dynamic way, and reduce redundancy either automatically or in a supervised yet user-friendly way. We first show with two study cases that *matrix-clustering* simplifies the interpretation of motif discovery results, and that a dynamic view of aligned logos can reveal biologically relevant motif variants. We then consider two applications encompassing complete databases, which show that the program regroups motifs bound by transcription factors of the same family, and can be used to explore the complementarity between multiple motif collections. This approach paves the way towards creating systematic non-redundant motif collections.

## MATERIAL AND METHODS

### Overview

*matrix-clustering* first computes a matrix of similarity between each pair of input PSSMs, runs hierarchical clustering to build a complete motif tree, which is then partitioned to generate motif clusters (Figure 1), based on a combination of thresholds on one or several motif similarity metrics. Within each cluster, PSSMs are then aligned. The results are displayed on a dynamic user-friendly web report enabling to collapse or expand subtrees at will.

**Figure 1.**
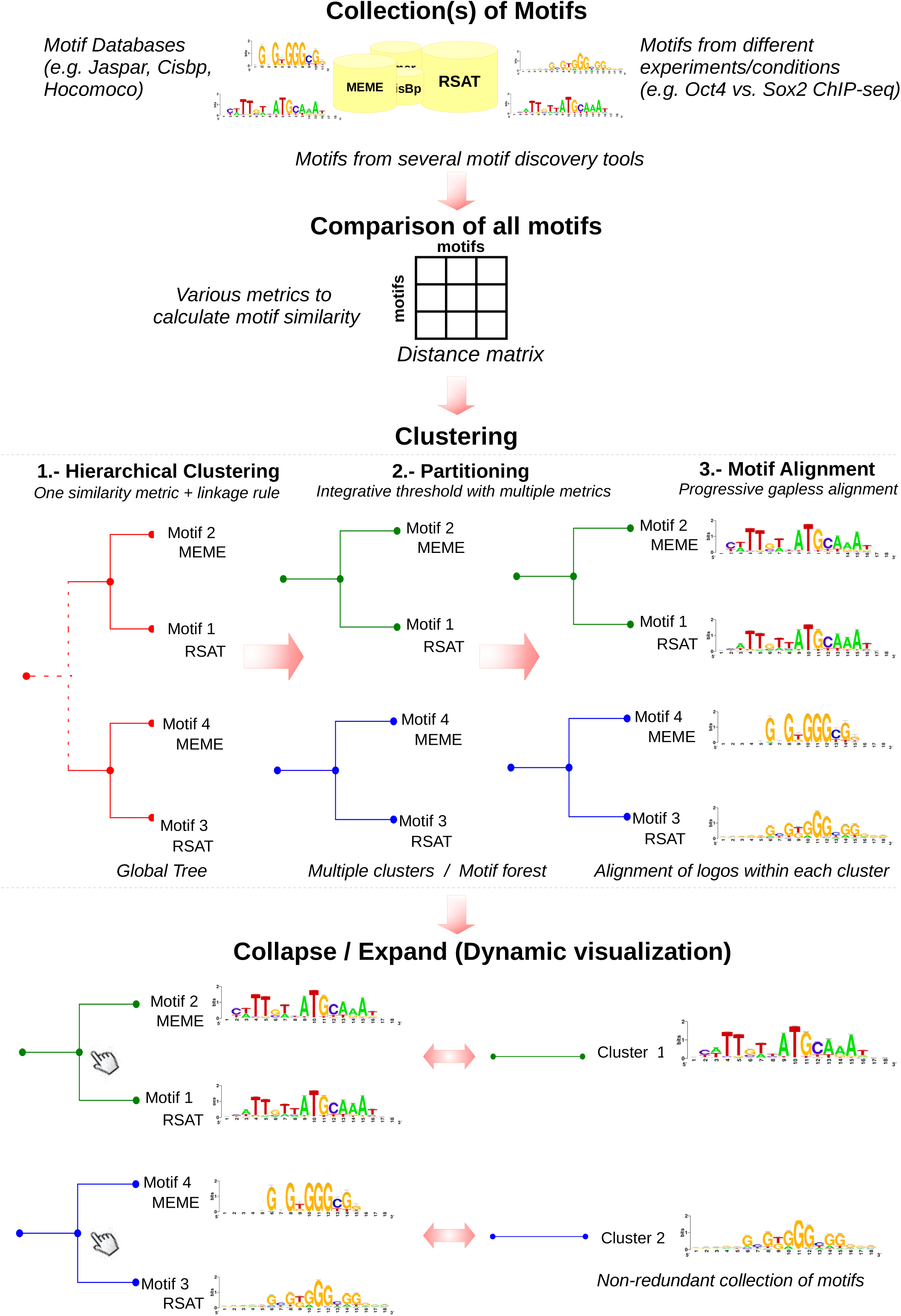
Schematic flow chart of the *matrix-clustering* algorithm. The program takes as input one (or several) collection(s) of PSSMs, and calculates the motif similarity using several metrics. One of these metrics is used to group the motifs with hierarchical clustering. A threshold consisting in a combination of metrics is used to cut the global tree in a set of subtrees. Each resulting tree then serves as a guide to progressively align the PSSMs. The PSSMs at the root of each tree are exported as non-redundant motifs. The trees can be collapsed or expanded at each node dynamically on the resulting Web page.

### Input formats and processing time

*matrix-clustering* receives as input one or several collections of PSSMs (provided as separate files) with an associated “collection name” (e.g. several PSSM collections obtained from different analyses or databases). This program supports different file formats: TRANSFAC (default), MEME, HOMER, JASPAR, etc., and has no restriction on the number of input PSSMs, but users should be aware that the processing time increases drastically with the number of motifs (Supplementary Figure 1). For small collections of motifs, the running time enables *matrix-clustering* usage via the website (e.g. 7 minutes for the first study case with 66 motifs). Large datasets can be treated with a stand-alone installation of the RSAT suite.

### PSSM comparison

Similarity between each pair of input PSSMs is calculated with the RSAT tool *compare-matrices (38, 39)*, which can compute multiple similarity metrics in a single run: Pearson correlation (cor), Sum Of Squared Distances (SSD), Mutual Information, Information correlation (Icor), Euclidean Distances (dEucl), Sandelin-Wasserman Similarity (SW), as well as width-normalized versions of some metrics obtained by dividing the total length of the alignment by the number of columns where the two PSSMs overlap: normalized correlation (Ncor), normalized information content correlation (NIcor), normalized Euclidian distance (NdEucl) (see Supplementary Notes for details). Each possible offset is tested for each pair of PSSMs in both orientations, and the program returns the best matching alignment.

### Hierarchical clustering

To build the *global hierarchical tree* encompassing all input PSSMs, the user must select one motif similarity metric (to make the motif-to-motif distance matrix) and one linkage method (average, complete or single). Some metrics directly measure distances (Euclidean, SSD, SW); for the metrics measuring similarities (e.g. cor, with a range from −1 to +1), the values are first transformed into dissimilarities (i.e. *Dcor = 2 − r*, where r is the correlation coefficient).

### Identification of motif clusters by tree partitioning

As the RSAT program *compare-matrices* (38) can return several metrics simultaneously, any combination of these can be selected to define thresholds for the partitioning step, thereby enabling to combine their respective advantages. The global tree is traversed in a bottom-up way and for each intermediate node, the selected metrics values are computed from all descendent leaves according to the chosen linkage rule (single, average, complete). Whenever an intermediate node fails to satisfy any of the threshold values, a new cluster is created by separating its two children branches.

### Progressive alignment of the PSSMs

Once the *global tree* is partitioned, each subtree is used as a guide to progressively align the PSSMs. They are first orientated (direct or reverse) and then shifted relative to each other. Note that this algorithm does not integrate internal gaps. This process produces one multiple alignment for each internal node of each tree, ending with a root alignment that encompasses all the PSSMs of a cluster.

### Branch-wise PSSMs, logos and consensus sequences

Once the PSSMs of each subtree have been aligned, *matrix-clustering* calculates for each node a branch-wise PSSM by summing (default) or averaging the frequencies of the descendent aligned motifs. It then generates the corresponding consensus sequences and logos. Branch-wise PSSMs introduced here are a generalization of the so-called familial binding profiles (FBP) (37).

### Dynamic visualisation of the clusters

The clusters are displayed as a PSSM forest, i.e. a collection of trees (one per cluster) with a logo at each leave. A unique feature of *matrix-clustering* is that trees can be browsed dynamically: each branch can be collapsed by clicking, and the resulting sub-tree is replaced by the logo of the branch PSSM, thereby enabling to produce customized motif trees (Figure 1).

### Cross-coverage of motif collections

When two or more motif collections are given as input, the cross-coverage indicates the percentage of the PSSMs from one collection that co-occur in clusters with PSSM from another collection. The cross-coverage of collection *A* by collection *B* (*c_A,B_*) is the number of PSSMs from *A* co-clustered with PSSMs from *B* ( |A_*with B*_|), divided by the total number of motifs in *A*(|*A*|).

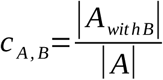

Reciprocally, the cross-coverage of collection *B* by collection *A* is computed as follows.

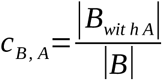

This asymmetrical comparison provides a more realistic interpretation of the importance of the intersection relative to the respective sizes of collections (e.g. a comparison between smaller and bigger databases). The cross-coverage is displayed as a heatmap, and a Venn diagram is drawn for each pair of collections. The percentage of motifs specific to each collection is also indicated.

### PSSM datasets of the study cases

Study cases 1 and 2: in order to illustrate the clustering of *ab initio* discovered motifs, we used 359 PSSMs obtained with the RSAT tool *peak-motifs* (15, 40) in 12 TF ChIP-seq peak-sets obtained from Chen et al (41). We also collected the PSSMs obtained by analysing one ChIP-seq peak set with MEME-ChIP (16) and Homer (42).

Study cases 3 and 4: for full database clustering, we analysed 24 taxon-specific collections from 18 databases (Supplementary Table 1): vertebrates (JASPAR (7), HOCOMOCO mouse and human (11), Cis-BP (9), Jolma 2013 “HumanTF” (4), Jolma 2015 “HumanTF_dimers” (13), Uniprobe (43), Fantom5 'novel' motifs (44), hPDI (45), epigram (46), Homer (42), Encode (47)), plants (JASPAR, Athamap (48), Cis-BP, ArabidopsisPBM (49) and Cistrome (14)) and insects (OntheFly (50), JASPAR, dmmpmm and idmmpmm (51), Cis-BP (9), FlyFactorSurvey (52), DrosphilaTF (53)).

### Availability

The tool *matrix-clustering* is freely available on the RSAT Web servers (http://www.rsat.eu/) (39). It can also be downloaded with the stand-alone RSAT distribution to be used on the Unix shell, allowing its inclusion in automated pipelines.

The complete results of the study cases are available on the supporting website: http://teaching.rsat.eu/data/published_data/Castro_2016_matrix-clustering/

### Implementation

*matrix-clustering* is implemented in Perl and R. The Logo trees are implemented in HTML5 with the D3 (54) JavaScript library for manipulating documents based on data (http://d3js.org/). The website dynamic elements are implemented using the JavaScript libraries Jquery (http://jquery.com/) and DataTables (http://www.datatables.net/).

## RESULTS

We have developed *matrix-clustering* to deal with the increasing number of motifs and reduce the inherent redundancy within collections. It takes as input one or more collections of PSSMs, measures the similarity between them using several motif comparison metrics, builds a similarity tree by hierarchical clustering, splits the initial tree to obtain one separate tree per cluster, generate a consensus and a logo for each branch of each tree, computes branch-wise PSSMs, and generates different graphical representations, including a dynamic visualization enabling flexible customization of the display (Figure 1).

### Choice of the default clustering parameters

Parameters of *matrix-clustering* were chosen based on a detailed comparison between clusters of 374 PSSMs from HOCOMOCO human TFBMs (11) and their classification in 21 families taken from the TFClass database (55). We tested four alternative similarity metrics (cor: correlation, Ncor: normalized correlation, Icor: information correlation, and NIcor: normalized information correlation), three linkage rules (single, average or complete), incremental series of partitioning threshold values on each metric (by step of 0.05), as well as combined thresholds applied on a metric and its normalized version (Ncor + cor, or NIcor + Icor). Based on this study, we defined the default parameters: the motif-to-motif similarity matrix is computed with the Ncor, with a minimal alignment width of 5 columns, the motif tree is built with the average linkage rule, and the partitioning threshold combine cor ≥ 0.6 and Ncor ≥ 0.4. The detailed results of the systematic evaluation, as well as the parameters used for each program, are described in the Supplementary Notes.

### Study case 1: identification of TF binding motif variants within motifs discovered with multiple tools in ChIP-seq datasets

It is common practice to perform *ab initio* motif discovery with several algorithms and to consider the motifs found by several approaches as robust predictions. Yet, some motif variants can be found only by a particular algorithm. This first study case aims at comparing motifs detected in ChIP-seq peaks with three motif discovery tools: RSAT *peak-motifs*, Homer and MEME-ChIP. We re-analysed the ChIP-seq peaks for the TF Oct4 (also named Pou5f1) in mouse embryonic stem cells (ESC) from Chen et al (41).

Altogether, the three tools produced 66 motifs: 22 discovered by RSAT *peak-motifs*, 25 by MEME-ChIP and 19 by Homer. *matrix-clustering* separated these 66 PSSMs into 13 clusters (Supporting website). The largest cluster regroups 37 PSSMs corresponding to Sox, Oct and other Oct-like motifs (Figure 2A). Since the name of the source collection is automatically displayed besides each logo (RSAT, MEME-ChIP, HOMER), we readily identify the robust motifs discovered by multiple tools, as well as motif variants detected by a single algorithm.

**Figure 2.**
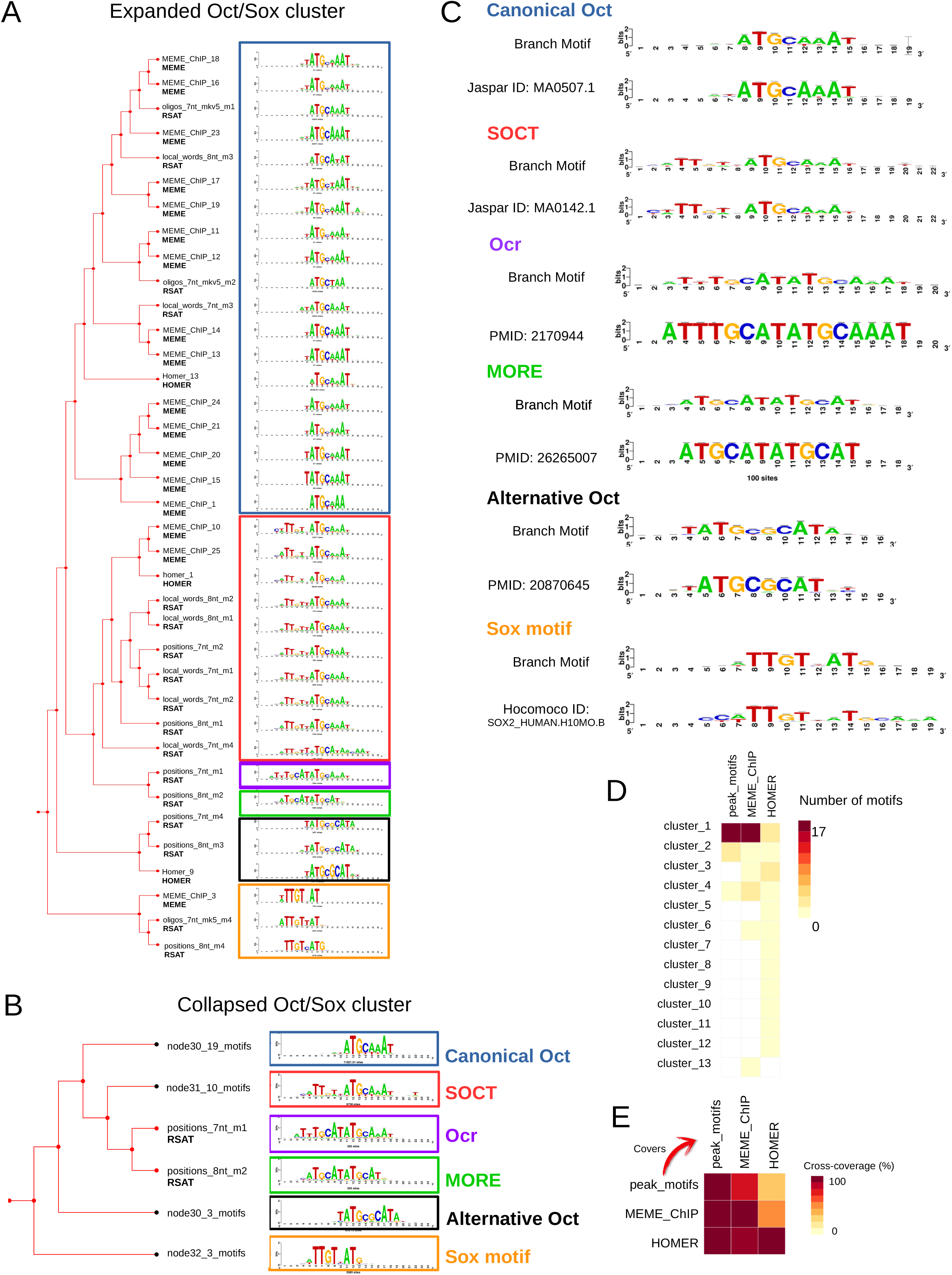
Clustering of PSSMs discovered in the Oct4 ChIP-seq peaks using several motif discovery tools. The TF peaks of Oct4 identified by Chen et al (41) were submitted to three *de novo* motif discovery programs: RSAT *peak-motifs*, MEME-ChIP and HOMER. All discovered PSSMs were clustered simultaneously by *matrix-clustering.* **(A)** Hierarchical tree corresponding to cluster_1 (37 motifs), where different Oct motif variants and Sox2 motifs are highlighted with different colored boxes. The leaves are annotated with the name of the submitted motif, and the name of its collection (one of the three programs). **(B)** Reduced tree showing six non-redundant motifs, obtained after manual curation of the cluster_1, by collapsing the branches. **(C)** Annotation of the six non-redundant variants (“branch PSSMs”) based on alignments to reference motifs (see main text). When available in databases (JASPAR or HOCOMOCO), the ID of the reference motif is indicated. Otherwise, it is replaced by the PMID of the publication mentioning the motif. **(D)** Heatmap summarising the number of motifs from each collection found in each cluster. **(E)** Heatmap of the cross-coverage between each collection.

We manually collapsed the cluster tree and identified six non-redundant motifs (Figure 2B) for which we searched for similarities in JASPAR vertebrates and HOCOMOCO Human (Figure 2C).These six motifs correspond to the canonical Oct4 (blue box on Fig. 2A and 2B), Sox2 (orange), the composite SOCT (Sox+Oct) motif (red) (56), an alternative configuration of Oct4 (black) (57), a palindromic Oct homodimer (More Palindromic Oct factor Recognition Element, MORE) (purple) (58), and an octamer-repeat (Ocr) (59). Of note, these last two motifs were only found by RSAT *peak-motifs* (Figure 2B).

The contributions of the respective motif discovery tools to the clusters are unbalanced. While RSAT *peak-motifs* contributes to three clusters shared with MEME and HOMER, MEME-ChIP raised one single-PSSM cluster (singleton) and HOMER six (Figure 2D). The cross-coverage between the tools (Figure 2E) confirms that *peak-motifs* and MEME show high overlap, whereas the HOMER motifs are quite dissimilar from those obtained with the other tools. Of note, many PSSMs found by HOMER only are actually of low-complexity (2-residue repeats) and are not likely to correspond to b*ona fide* TFBMs.

Altogether, this study case demonstrates that *matrix-clustering* can guide and accelerate human-based reduction of a highly redundant collection of motifs, produced by running several motif discovery tools on the same sequence set. The clustering moreover highlights the existence of TFBM variants and combinations (e.g. homodimers, heterodimers).

### Study case 2: identification of exclusive or shared motifs between various ChIP-seq experiments

We extended our previous analysis to the 12 TFs studied by Chen at al (41) in order to identify common and set-specific motifs among the ChIP-seq peak sets. We ran RSAT *peak-motifs* in each peak set separately and obtained 359 PSSMs, regrouped by *matrix-clustering* into 28 clusters (Supporting website).

Some clusters contain set-specific motifs, e.g. Stat3 (cluster_12), Nanog (cluster_14), Ctcf (cluster_17) and Zfx (cluster_18) (Figure 3A). Other clusters contain motifs found in two or more peak sets: the Sox (cluster_10), Myc (cluster_5) and Oct motifs (cluster_1) are respectively found in three (Oct4, Sox2, and Nanog), two (nMyc and CMyc) and six (Oct4, Sox2, Nanog, Stat3, Tcfcp2l1, cMyc) peak sets (Figure 3A). These TFs are known to cooperatively regulate common target genes, explaining why their motifs are found across multiple peak sets (41, 56). The cross-coverage heatmap (Figure 3B) provides a global view of the content similarity between motif collections. This representation confirms that PSSMs discovered in Oct, Sox and Nanog peak sets are highly similar, consistent with the fact that these TFs co-occur in shared enhancers (41). This is also the case for the cMyc and nMyc motifs, as well as for E2f1 and Zfx, which are functionally related as histone genes regulators (60). By contrast, the motifs discovered in CTCF peak sets are mostly specific to this collection. This study case shows that handling multiple motif collections (feature unique to *matrix-clustering*) can highlight their similarities and differences.

**Figure 3.**
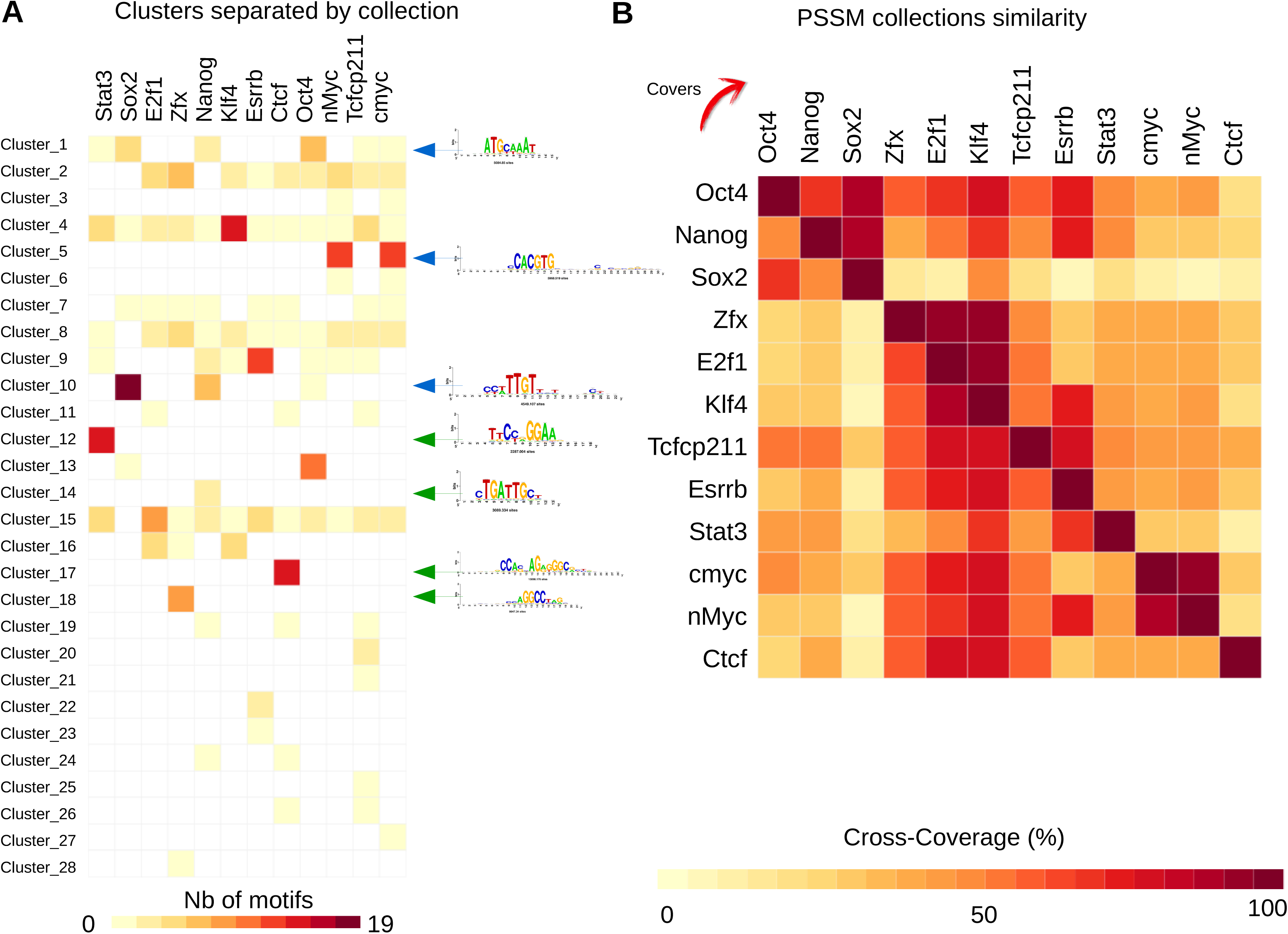
Clustering of 12 sets of PSSMs discovered in mouse ESC TF ChIP-seq peaks. **(A)**Matrix showing the cluster composition by motif collection. Examples of motifs found in one or several collections (and their corresponding logos) are indicated with green and blue arrows, respectively. **(B)** Heatmap showing the cross-coverage between the 12 motif collections corresponding to the ESC TF peak-sets.

### Study case 3: Complete database analyses highlights relationships between motif clusters and TF families

We evaluated whether a clustering of complete motif databases enables (i) to identify redundancy between motifs, and (ii) to regroup PSSMs from the same TF family. TFs are classified in families according to their DNA-binding domains (DBD) (55, 61), which usually recognize similar binding sites. TF belonging to the same families are thus often associated with similar TFBMs, which constitute a source of redundancy.

We clustered the complete set of taxon-specific motifs from JASPAR (vertebrates and insects), and species-specific motifs from HOCOMOCO (human and mouse). The clustering of JASPAR insects (133 motifs) reveals a large cluster of 70 PSSMss (Figure 4A; Supporting website) encompassing almost half of the database. This corresponds to homeodomain-containing TFs, whose binding motifs are characterized by the core consensus 5'-TAAT-3' (62). The dynamic browsing capabilities of *matrix-clustering* enable to manually reduce these 70 PSSMs to 10 distinct motifs (Figure 4B). The numerous members of this family in the insect database reflect an annotation bias, as most of these PSSMs result from a single analysis covering many homeodomain TFs (63).

By contrast in vertebrates, the 641 human PSSMs of HOCOMOCO are reduced to 127 small clusters (Figure 4C). We obtained similar results for JASPAR vertebrates and HOCOMOCO mouse collections (Supplementary Figures 2A and 2B, supporting website). As HOCOMOCO includes the information about TF families imported from TFclass (55), we analysed the correspondence between clusters produced by *matrix-clustering* and these TF families. The majority of the clusters (77 out of 127) indeed regroup motifs bound by TFs from a single family (Figure 4D). Furthermore, most of the other clusters actually regroup TFs belonging to different families of the same class. The remaining clusters encompass TFs from different classes but nevertheless bound to similar motifs, and thus correctly grouped by *matrix-clustering.*

**Figure 4.**
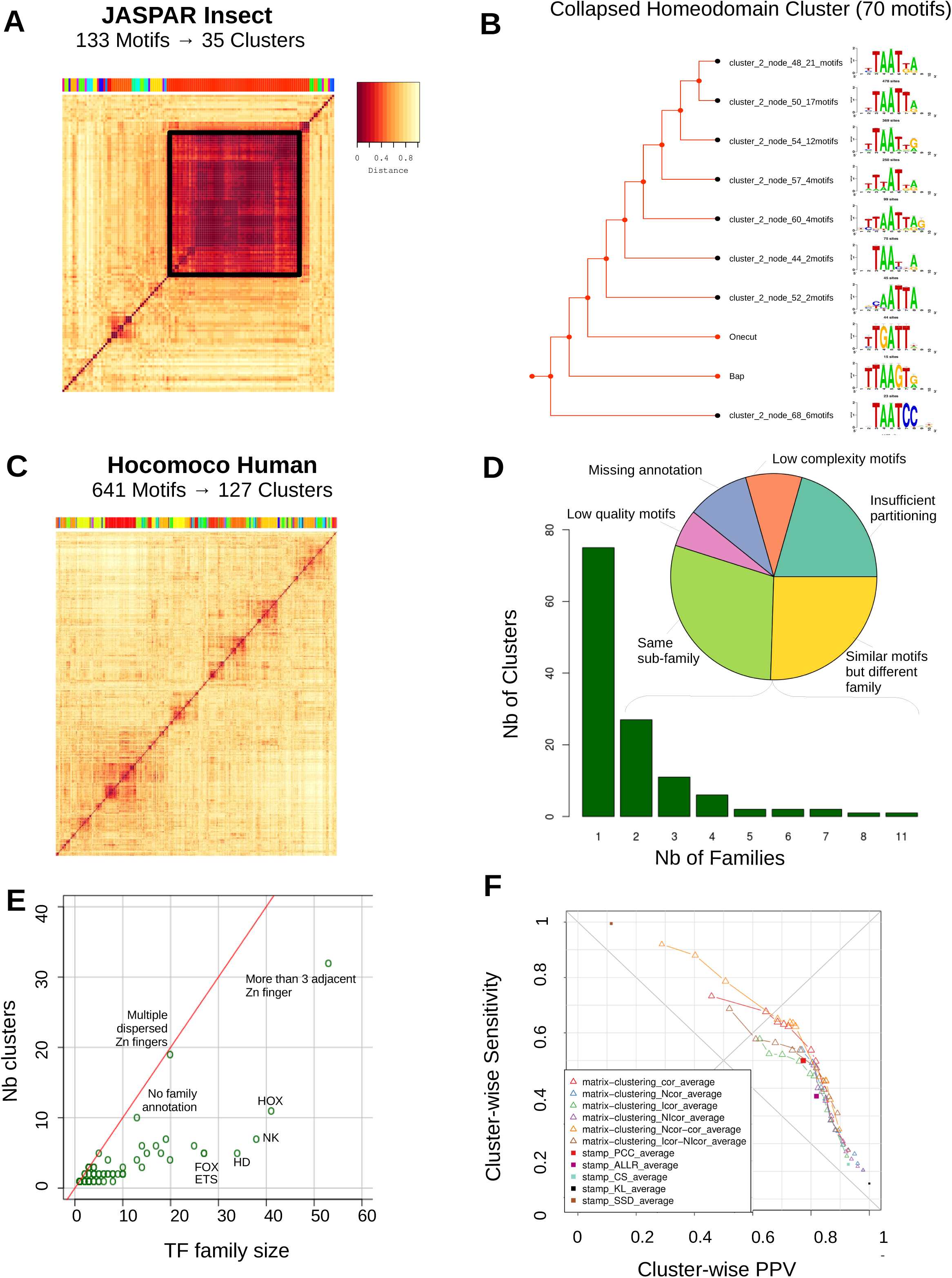
Clustering of complete Insect and Human motif databases. **(A)** Heatmap representing the similarity (Ncor) between all 133 PSSMs of JASPAR Insects. The 40 clusters found are indicated with a colored bar above the heatmap. The black square emphasizes the large cluster (almost half of the PSSMs) containing the very similar Homeodomain motifs. **(B)** The 70 Homeodomain motifs were manually reduced by collapsing the tree branches into ten motifs. The collapsed tree is displayed along with the corresponding aligned branch motifs. **(C)** Heatmap representing the similarity (Ncor) between all 641 PSSMs of HOCOMOCO Human. **(D)** Repartition of the clusters formed from HOCOMOCO Human with TF families. The bar plot indicates that most clusters are composed of a single TF family. The pie chart illustrates the reasons for observing multiple TF families in a single cluster. **(E)** Scatterplot comparing the number of members of each TF family as a function of the number of covered clusters. The name of the families with more than 20 members are shown. **(F)** Scatterplot showing the trade-off between sensitivity and specificity by clustering PSSMs from the same family with either *matrix-clustering* or STAMP, using different parameters to compute a similarities between each pair of input matrices, build the trees and define the clusters. For *matrix-clustering*, the curves denote a series of tests performed with different threshold values on the same dissimilarity metric. For STAMP the number of clusters is defined automatically. Dot sizes are proportional to the geometric accuracy. The ideal clustering would be in the top-right corner.

Reciprocally, for each TF family we counted the number of covered clusters (Figure 4E, Supplementary Figure 3). Among the 78 families from HOCOMOCO, 29 are consistently packed in a single cluster, 10 in two clusters, and 16 in three clusters. On the other extreme, some TF families are split into many clusters, in particular the Zinc finger families (e.g. for the family “Factors with multiple dispersed zinc fingers”, each PSSM comes as a separate cluster). This dispersion is perfectly consistent with the well-known properties of these TFs: the sequence bound by each Zinc finger domain is determined by the four specific amino acids entering in contact with the DNA (64).

As above mentioned, we explored the impact of clustering parameters on the correspondence between clusters of PSSMs from Human HOCOMOCO (11) and the families of the bound TFs (see section “*Choice of the default parameters*” and Supplementary Notes). The highest accuracy was achieved with *Ncor* as matrix-to-matrix comparison metric, a tree built with the average linkage rule, which is partitioned according to a combined threshold on *Ncor* (≥ 0.4) and *cor* (≥ 0.6) (Figure 4F).

This study case demonstrates how *matrix-clustering* can handle large collections of PSSMs and automatically reduce their redundancy within a database, while correctly regrouping motifs belonging to the same TF Family.

### Study case 4: Comparison and integration of multiple motif databases

To evaluate inter-database redundancy and to automatically produce a non-redundant motif set, we clustered 24 motif collections and measured their cross-coverage (see Supplementary Table 1 and Material and Methods for the complete list of collections).

We first merged these public databases to obtain three taxon-specific collections for insects (7 databases; 1895 PSSMs), plants (5 databases; 1590 PSSMs) and vertebrates (12 databases; 7781 PSSMs), respectively. We then applied *matrix-clustering* and obtained 354 clusters for insects (19% of the total merged PSSM collection), 306 for plants (19%) and 1757 for vertebrates (33%) (supporting website). In order to obtain non-redundant motifs whilst preserving specificity, we used more stringent partitioning criteria than the default (cor >= 0.8 and Ncor >= 0.65): the threshold on correlation ensures that the clustered motifs are highly similar and the additional threshold on normalized correlation selects the alignments covering most of the motif lengths, in order to separate composite motifs (e.g. bound by a TF dimer) from their elementary components.

We then explored the mutual overlap between the original collections by computing the cross-coverage (Figure 5). For the insect databases, Cis-BP, OnTheFly, FlyFactorSurvey and JASPAR are the most similar to each other, while DrosophilaTF is drastically different from all of them (Figure 5A), likely because this collection was built by selecting motifs discovered exclusively on Drosophila promoters, and whose binding factors are unknown (53).

**Figure 5.**
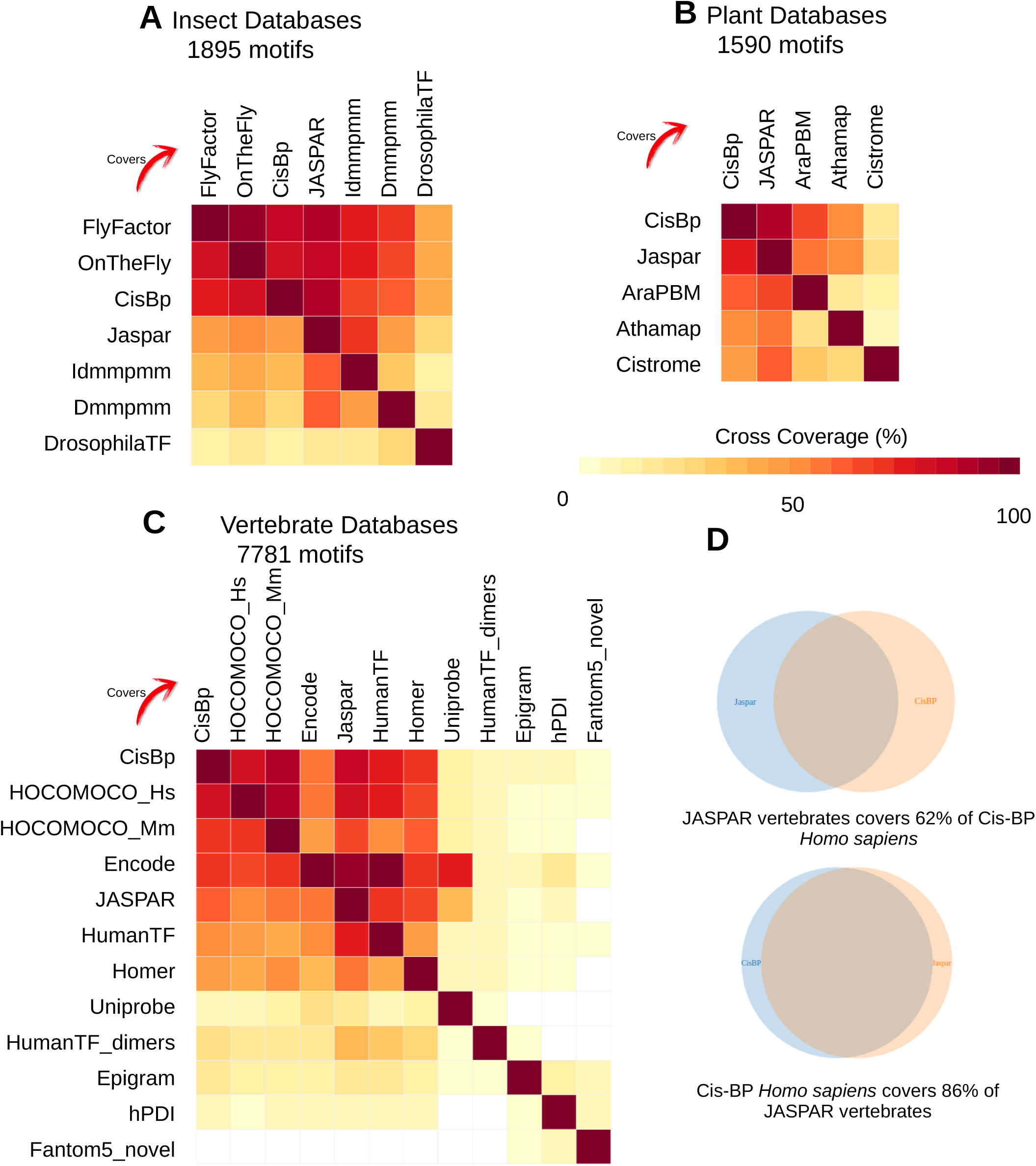
Cross-coverage of public motif databases. Several full public collections were merged and clustered, separately by taxa. The heatmaps of the cross-coverage between each collection is plotted for **(A)** seven insect collections, **(B)** five plant databases, and **(C)** twelve vertebrate databases. The heatmaps show the cross-coverages for each pair of databases. Note that the heatmaps are not symmetrical because the numbers of motifs in the different databases differ. **(D)** Venn diagrams showing the asymmetry of cross-coverage between two databases with different sizes.

For the plant databases, JASPAR and Cis-BP are most similar to each other (Figure 5B), which is coherent with Cis-BP being an integrative motif collection encompassing other public collections (including JASPAR). The three other databases focus on sets of motifs characterized by specific experimental methods (PBM for ArabidopsisPBM, binding sites curated from literature for Athamap, DAP-seq for CisTrome).

Regarding vertebrates, five databases have a similar content (HOCOMOCO human and mouse, JASPAR, Cis-BP, Jolma 2013 “HumanTF”), which is explained by the integration of HOCOMOCO and JASPAR in Cis-BP, as well as by the similarity of the original datasets used to build the TFBMs (mostly public ChIP-seq, Selex-seq and PBM), yet with different algorithms (Figure 5C). Note that the cross-coverage is not reciprocal since the number of motifs and the motif diversity differ among these databases. For example JASPAR includes 62% of the content of Cis-BP, whereas the latter encompasses 86% of JASPAR motifs (Figure 5D). We observed that the motif diversity is not proportional to the database size (e.g. the 641 JASPAR vertebrate PSSMs cover 82% of the 1800 Cis-BP Human PSSMs). In contrast, the contents of the remaining databases differ considerably according to the different methods and data used to build the motifs: a single type of data (Uniprobe, derived from PBMs only), restricted numbers of sites (hPDI, 17 sequences per motif on average), data from ChIP-seq experiments targeting histone marks in different cell types (epigram), or motifs modelling TF dimers (HumanTF_dimers). The low cross-coverage of Fantom5 collection of “novel” motifs is consistent with th definition if this database, which is restricted to motifs without any matches in reference databases (44).

In summary, this study case highlights how *matrix-clustering* can be used to automatically reduce motif redundancy across multiple databases into non-redundant taxon-wise motif collections (available as Supporting files 1-3 and on the supporting website) encompassing several thousands of PSSMs. The concise representation provided by the cross-coverage heatmap enables to intuitively grasp the overlap between each pair of individual collections.

### Comparison with alternative motif clustering tools

RSAT matrix-clustering is the only tool supporting dynamic browsing of motif trees with custom collapse/expansion of branches, and providing multiple ways to inspect the results: motif forest with branch motifs at each level of each tree, similarity heatmap, searchable table of motifs and clusters, comparison between multiple collections with contingency tables summarizing relationships between clusters and collections, as well as cross-coverage between collections. See Table 1 with a list of features supported by existing motif clustering tools. This flexibility has a cost in computing time (see Supplementary notes for a comparison of time efficiency between STAMP and matrix-clustering).

We performed a detailed comparison with STAMP, varying its parameters, and observed that its accuracy (based on a single metric) is lower than *matrix-clustering* using two metrics to separate the clusters (Figure 4F, see details in Supplementary Notes).

We furthermore submitted two of our following study cases to several motif clustering tools (using default parameters): STAMP (22), m2match (25), Matlign (26) and Gmacs (28). This analysis was restricted to case studies 1 and 3, since no other tool currently supports the clustering of multiple collections. The results are detailed in the Supplementary Notes.

## DISCUSSION

With the advent of large-scale experimental approaches to uncover TF binding specificity such as ChIP-seq, Selex-seq and PBMs, the number of TFBMs has recently exploded, and motif redundancy is becoming a critical bottleneck for sequence analyses. Although many software tools are available to measure motif similarity, only a few tools are truly specialized in motif clustering. A basic survey of motif clustering tools and their functionalities (Table 1) revealed many limitations that prompted us to develop *matrix-clustering.*

A key feature that distinguishes *matrix-clustering* from the other tools is its dynamic interface to browse clustered PSSMs. This feature substantially facilitates the manual control of cluster visualization and reduces the time for human analysis of motif sets. Notably, this visualization has enabled us to identify the Ocr motif in the Oct4 ChIP-seq peaks (Figure 2). This motif was already present in our previous analysis of the same dataset (15), but we had not been able to detect this subtle variation among all other unclustered motifs. We thus expect that this dynamic visualisation of motif clusters will be beneficial to both experts and non-experts users. Furthermore, *matrix-clustering* dynamic interface can be used and integrated in the website of motif databases.

Our method relies on hierarchical clustering with a bottom-up partitioning. The tree is thus segmented based on the similarity between all the descendant PSSMs of each branch, which strongly differs from the usual cut-off at an arbitrary height of the clustering tree. We evaluated an alternative segmentation method called *dynamic tree cut*, which relies on tree topology to produce balanced clusters (66), but we kept our approach because it allows to cut the tree based on motif similarity rather than on the sole tree topology. One caveat of hierarchical clustering is to produce 'frozen' clusters, i.e. nodes regrouped early in the tree cannot be relocated in later steps (28). Note that some motif clustering tools avoid this problem by using iterative assignment algorithms, such as k-medoids (28), and that STAMP circumvents it by refining the tree a posteriori (22).

Partitioning thresholds should be tuned to reach the desired granularity of clusters. Based on the systematic evaluation of HOCOMOCO motifs we used as default thresholds (Ncor >= 0.4 and cor >= 0.6) to group the TF binding variants and motifs from the same TF family within the same cluster. However, in order to favour specificity and obtain non-redundant collection of motifs (study case 4), stringent thresholds can be used (Ncor >= 0.65 and cor >= 0.80).

Several databases like JASPAR and HOCOMOCO already provide non-redundant collections, obtained by a time-consuming manual curation, which will become complicated to maintain with the increasing number of motifs. Of note, in motif databases, the term non-redundant denotes the restriction to one PSSM per TF (7). However, distinct TFs may also bind very similar motifs (e.g. Oct4, Oct9, and Oct11), and in some cases a same TF might bind to alternative motifs (e.g. TF complexes, or multi-domain TFs). In this study, the term non-redundant refers to a single PSSM summarizing a set of highly similar motifs, independently of the binding TF.

Reducing the size of motif collections is becoming crucial to limit the processing time of tools relying on full motif databases (e.g. motif enrichment, motif comparisons, identification of regulatory variants). As a proof-of-concept, we have shown that *matrix-clustering* can be used to compare full collections, but also to drastically reduce inter-database redundancy: in case study 4 we produced non-redundant motifs collections that reduced the insect, plant and vertebrate collections to 19%, 19% and 32% of their original sizes, respectively. We thus expect that meta-databases, such as footprintDB (10) or Cis-BP (9) could benefit from *matrix-clustering* to offer non-redundant motif collections.

Non-redundant motif collections would reduce computing time when scanning big sequence sets with large collections of PSSMs. However, it should be noted that merged motifs resulting from clustering are by definition less specific than the original motifs, more so if they have a poor quality. Still, for motifs built from a few binding sites, a merged motif could be more specific (Supplementary Notes). We suggest that merged PSSMs could be used to represent a group of similar motifs to reduce computing time for tasks affected by motif redundancy (e.g. comparison of discovered motifs with reference databases). For more precise tasks, such as TFBS prediction, they can be suboptimal.

The possibility to cluster several collections simultaneously makes *matrix-clustering* a versatile tool, as demonstrated with the four case studies considered (identification of motif variants, integration of motifs found by multiple motif discovery tools, comparison of motifs obtained from many collections). The same tool could be used to compare motifs obtained in different experimental conditions. Given the compatibility with many PSSMs formats (TRANSFAC, MEME, HOMER) and its Web access, this tool will be of interest to the broad community of biologists and bioinformaticians involved in the analysis of regulatory sequences.

## ACKNOWLEDGEMENT

We wish to thank to Bruno Contreras-Moreira, Aïtor Gonzales, Carl Herrmann, Samuel Collombet, Roberto Tirado-Magallanes, Lambert Moyon and Coby Viner for suggestions to improve the method and the visualization. We are also thankful to the JASPAR team for check and validate the clustering of JASPAR motif collections.

